# A crosswalk to align the GSS and NIH grants databases based on university names

**DOI:** 10.1101/2021.02.25.432534

**Authors:** Christopher L. Pickett

## Abstract

Notable reports over the past dozen years have recommended the federal government and others improve data collection on the research workforce in the United States. The federal government already collects a wealth of data, but important datapoints, like the number of biomedical postdocs working in the U.S., for example, are still not well defined. Furthermore, of the data that are collected, differences in collection method and data naming conventions, like inconsistent naming of universities, hinders our ability to merge information across different datasets. Here I describe the creation of three macros meant to align the National Center for Science and Engineering Statistics’ Survey of Graduate Students and Postdoctorates in Science and Engineering (GSS) with NIH grants databases based on consolidating university names under a single name common to both databases. Aligning these databases will allow for a deeper understanding of how various federal and university policies affect the number trainees, grants, and funding at individual institutions.

## Introduction

Collecting and analyzing data on the scientific workforce is necessary to develop policies that ensure an equitable, vibrant research enterprise. The federal government supports many data collection efforts and databases that are invaluable sources of information on the research funding and the workforce. For example, the National Science Foundation’s (NSF) National Center for Science and Engineering Statistics (NCSES) coordinates and conducts a series of surveys of universities, scientists and engineers, and trainees to provide snapshots of the science and engineering workforce (https://www.nsf.gov/statistics/). And the National Institutes of Health publishes a wealth of information on each grant the agency awarded since 1985 (https://report.nih.gov/). In addition, non-federal databases also provide necessary information about the research workforce including those from the Association of American Medical Colleges (https://www.aamc.org/data-reports/faculty-institutions/faculty-roster) and the Coalition for Next Generation Life Sciences (http://nglscoalition.org).

Despite this volume of information, there are still dimensions to the research enterprise that are poorly understood. For example, we do not have a firm grasp on the number of postdocs working in biomedical research (*1-4*). For this reason and others, multiple reports on the research enterprise, and the biomedical research enterprise in particular, recommended the federal government improve data collection on the research workforce (*5*). Recently, universities began collecting and publishing the career outcomes of their Ph.D. and postdoc alumni (*6-9*). These efforts satisfy some of the recommendations from prior reports calling for more extensive data collection on the research workforce, but they do not solve the entire problem.

In addition to collecting more data, it is important to fully interrogate the data that have already been collected. One barrier to this is that not all federal databases are relatable. For example, the NIH publishes data on grant recipients including their institution, the type of grant and how much money was awarded. Similarly, the NCSES’s Survey of Graduate Students and Postdoctorates in Science and Engineering (GSS) publishes information on how many graduate students and postdocs are enrolled in specific disciplines at individual institutions. But while both databases list vital information on institutions, these databases do not present data in a way that we can readily understand how much funding an institution receives on a per trainee basis, for example. In the simplest example of how these databases do not align, the GSS lists the University of Utah as “University of Utah, The” while the NIH lists the institution as “UNIVERSITY OF UTAH.” These are clearly the same institution, but the computer programs needed to conduct these analyses will not recognize them as such because of differences in formatting. Furthermore, institutional affiliations assumed in one database but not the other, name changes, university mergers, and other differences in naming conventions complicate aligning the GSS and the NIH databases.

Here, I describe tools to link the NCSES’s GSS with the NIH grants database. I created Microsoft Excel macros that consolidate and unify the naming conventions and affiliations of universities within the GSS and NIH databases individually, and then again as a crosswalk between the two databases. By ensuring consistent naming, these two databases become relatable, allowing deeper investigations into the how the number of trainees at individual institutions is affected by changes in funding to that institution.

## Methods

National Institutes of Health grant information for 1985 through 2016 was downloaded from NIH ExPORTER (https://exporter.nih.gov/). The microdata for the Survey of Graduate Students and Postdoctorates in Science and Engineering (GSS) for 1985 through 2016 was downloaded from the National Center for Science and Engineering Statistics division of the National Science Foundation (https://nsf.gov/statistics/srvygradpostdoc/pub_data.cfm).

My intention was to relate trainee numbers at the institution level in the GSS dataset with grant awards and direct costs to institutions in the NIH dataset. The various unique codes assigned to institutions in either dataset were not the same, and the best way to connect the two datasets was based on institutional names. However, even within datasets, universities were not consistently named.

### GSS consolidation

In the GSS dataset between 1985 and 2016, I first isolated the “institution_id” and “Institution_Name” fields. The “institution_id” is a unique code assigned to institutions and is useful for identifying institutions that had multiple names over time. For example, institutional ID code 1064 is linked to California State University in Bakersfield, which is listed in the GSS as “California State Univ, Bakersfield” from 1985 to 2009, “California State University-Bakersfield” from 2010 to 2011, and “California State University, Bakersfield” from 2012 to 2016. It is not clear if the name change is due to changes at the institution level or at the level of the GSS. However, to understand changes in the trainee population of California State University in Bakersfield over time, these entries were consolidated.

Standard Microsoft Excel (Microsoft Excel for Mac V16.46; Microsoft Corp., Redmond, WA, USA) formulas and processes were used to identify institutions with the same ID but different names. Next, a visual scan of institution names was conducted to identify institutions that were the same but not identified in prior steps. This step was informed, in part, by previous work (*10*). A list of GSS institutions was arrayed vertically with groups of aliases for institutions (hereafter “associated names”) appearing horizontally next to the main institution name (hereafter “common name”) (Table S1).

Between 1985 and 2016, the GSS had 905 unique entries for institutions in the “Institution_Name” field. Of these, 176 institutions had multiple names in the GSS database: 1 institution had five names, 12 institutions had four names, 50 institutions had three names, 113 had two names. The remaining 476 had only a single name in the 30 years of the GSS sampled here. This left 652 unique common names in the GSS database (Table S1).

### NIH consolidation

To consolidate data within the NIH dataset, I started from the merged list of institutions published previously (*10*). Then, similar to the GSS data, I used standard Microsoft Excel functions to identify institutions with different “ORG_NAME” but the same “ORG_DUNS.” The “ORG_DUNS” value functions analogously to the “institution_id” in the GSS. I then conducted a visual scan of institution names combined with internet searches to determine the university affiliations of hospitals, research centers, and institutions, as well as to identify name changes and other rearrangements.

Because I wanted to align the NIH dataset with the GSS, I made the decision to consolidate information on hospitals, research centers, and other organizations based on the Ph.D.-granting institution they were affiliated with. The Dana-Farber Cancer Institute, for example, was consolidated under Harvard University. While the Dana-Farber receives NIH grants, the graduate students working in labs at the institute receive their degrees from Harvard University.

A list of NIH institutions was arrayed vertically with groups of associated names appearing horizontally next to the common name (Table S1). Between 1985 and 2016, the NIH distributed grants to 2,706 institutions. Of these, 206 institutions had multiple: 118 institutions had two names, 36 institutions had three names, and 52 institutions had four or more names. The remaining 1,943 institutions had only a single name in the ORG_NAME field in the datasets sampled here. This left 2,149 unique common names in the consolidated NIH database (Table S1).

### Crosswalk

To make a crosswalk between the NIH and GSS data, I first attempted to match NIH and GSS common names using standard Microsoft Excel functions. This met with limited success primarily because of simple differences in naming conventions. I next conducted a visual scan of the datasets to match institutions across them. The NIH and GSS institution arrays were then combined with groups of associated names for institutions appearing horizontally next to the common name (Table S1). Of the 652 entries in the consolidated GSS database, 375 entries (57.5%) had a corresponding entry in the NIH database.

### Macro creation

I chose to create macros in Microsoft Excel to accelerate the consolidation of university names in the GSS and NIH datasets. Because all of the data cited here are downloadable as Excel-readable spreadsheets, the macros could be applied to the spreadsheets prior to additional analyses or conversion to other programs.

Once I had identified all associated names for a university in the GSS dataset above, I created the NSF_University_update macro that searches for each instance of an associated name and replaces it with the common name (Table S2). When the macro is run on a GSS dataset with the “Institution_Name” column, all associated names for an institution are replaced with the institution’s common name, as defined in Table S1.

Similarly, once I had identified all associated names for a university in the NIH dataset, I created the NIH_university_update macro that searches for each instance of an associated name and replaces it with the common name (Table S3). When the macro is run on an NIH dataset, all associated names in the “ORG_NAME” field are replaced with the institution’s common name, as defined in Table S1.

I then created the NSF_NIH_university_crosswalk macro that searches for each instance of the common name used for the NIH dataset and replaces it with the common name used in the GSS dataset (Table S4). When the macro is run on an NIH dataset, all common names in the “ORG_NAME” field that have a match in the GSS dataset are replaced with the common name used in the GSS dataset. After running these three macros, all entries in NIH RePORTER and the GSS between 1985 to 2016 can be compared based on the institution’s common name.

### Isolating biological and biomedical trainees

The NIH primarily supports biological and biomedical scientists, while the GSS surveys the numbers and demographics of trainees from all science and engineering fields. To better understand the relationship between NIH funding and biological and biomedical trainees, I selected GSS data based on the gss_code field corresponding best to biological and biomedical trainees: all 600 series codes (biological and biomedical sciences) and code 950 (neurobiology and neuroscience). I also included all 700 series codes (health sciences) because research in these departments is largely funded by NIH grants, and the research done in these departments often overlaps with those in the biological and biomedical departments.

## Results

The NIH Data Book (https://report.nih.gov/nihdatabook/) and other resources from the NIH give some indication of trainee support, but these data are not broken down to the level of individual institutions. The GSS has information on the numbers and demographics of graduate students and postdocs at individual institutions, but no information on how much training support each institution receives. Applying the macros designed here to data from the NIH and the GSS make it possible to make direct comparisons of information from both datasets.

To test the macros, I applied them to the GSS and NIH databases, and then I determined the 10 institutions that reported the most graduate students and postdocs in 2016 via the GSS. These institutions were in descending order of number of trainees: Harvard University, Johns Hopkins University, the University of Washington, the University of Michigan, Stanford University, the University of North Carolina-Chapel Hill, the University of Minnesota, Columbia University, the University of Florida, and Boston University. They ranged from 1,793 to 7,061 trainees in 2016 as reported to the GSS (Table 1). The primary mechanisms of direct trainee support provided by the NIH are the F31, F32 and T32 grant awards. The ten institutions analyzed here received between 23 and 308 F31, F32, and T32 training awards from the NIH in 2016, totaling between $3 million and nearly $53 million (Table 1).

**Table 1:** Aligning the GSS and NIH databases allows for additional insights into biomedical workforce support.

| Institution<sup>a</sup> | Trainees<sup>b</sup> | Awards<sup>c</sup> | Funding<sup>c</sup> | Trainees/Award<sup>d</sup> | Funding/Trainee<sup>d</sup> |
| --- | --- | --- | --- | --- | --- |
| Harvard University (1/35) | 7061 | 308 | \$52,808,223 | 22.9 | \$7,479 |
| Johns Hopkins University (1/8) | 3437 | 144 | \$29,658,993 | 23.9 | \$8,629 |
| University of Washington (1/17) | 2409 | 121 | \$23,444,019 | 19.9 | \$9,732 |
| University of Michigan (1/2) | 2319 | 107 | \$20,544,426 | 21.7 | \$8,859 |
| Stanford University (1/4) | 2311 | 149 | \$22,514,774 | 15.5 | \$9,742 |
| University of North Carolina at Chapel Hill, The (3/3) | 2285 | 128 | \$18,902,137 | 17.9 | \$8,272 |
| University of Minnesota (1/6) | 1984 | 56 | \$10,432,064 | 35.4 | \$5,258 |
| Columbia University in the City of New York (4/9) | 1913 | 105 | \$17,397,834 | 18.2 | \$9,095 |
| University of Florida (1/1) | 1891 | 23 | \$3,063,741 | 82.2 | \$1,620 |
| Boston University (1/6) | 1793 | 42 | \$6,947,063 | 42.7 | \$3,875 |

<sup>a</sup>Common name used after consolidation with macros described here. In parentheses, the first number indicates the number of entries consolidated with the GSS dataset and the second number indicates the number of entries consolidated in the NIH dataset.
<sup>b</sup>The number of graduate students and postdocs at an institution in 2016, as derived from the GSS.
<sup>c</sup>The number of awards and amount of funding from F31, F32, and T32 awards received by institutions in 2016, as derived from the NIH dataset.
<sup>d</sup>The number of trainees per F31, F32, and T32 awards received in 2016, or the amount of F31, F32, and T32 funding received in 2016 for every trainee at that institution.

The consolidation of institutions described here allows for a more direct relation of data between the GSS and the NIH datasets and adds a new dimension of analysis. For example, these ten institutions ranged from 15.5 trainees per F31, F32, and T32 award (Stanford University) up to 82.2 trainees per award (University of Florida) (Table 1). These data indicate a wide disparity among these ten institutions in how many training-specific awards they received on a per trainee basis.

T32 awards can support multiple trainees. As such, the awards per trainee discussed above may not give a complete picture of trainee support at an institution. Instead, I analyzed the amount of F31, F32, and T32 funding each institution received per trainee. In 2016, the institutions analyzed here received between $1,620 per trainee (University of Florida) up to $9,742 per trainee (Stanford University; Table 1). These data give a better sense of the training-specific resources individual institutions received in 2016.

Additionally, all research institutions supporting graduate students or postdocs can be visualized with these metrics. Despite the seeming large differences in trainees per training award, the variance across the research enterprise is much greater than that observed in the ten institutions analyzed above. Specifically, institutions that have fewer training awards tend to have more trainees per award and are more likely to have starkly different numbers of trainees per award relative to institutions that have the same number of awards (Fig. 1A). Similarly, the ten institutions analyzed above received more training money per trainee than most of the research enterprise (Fig. 1B). These data indicate that, despite training more graduate students and postdocs, these institutions draw training dollars well above the average institution.

**Figure 1:**
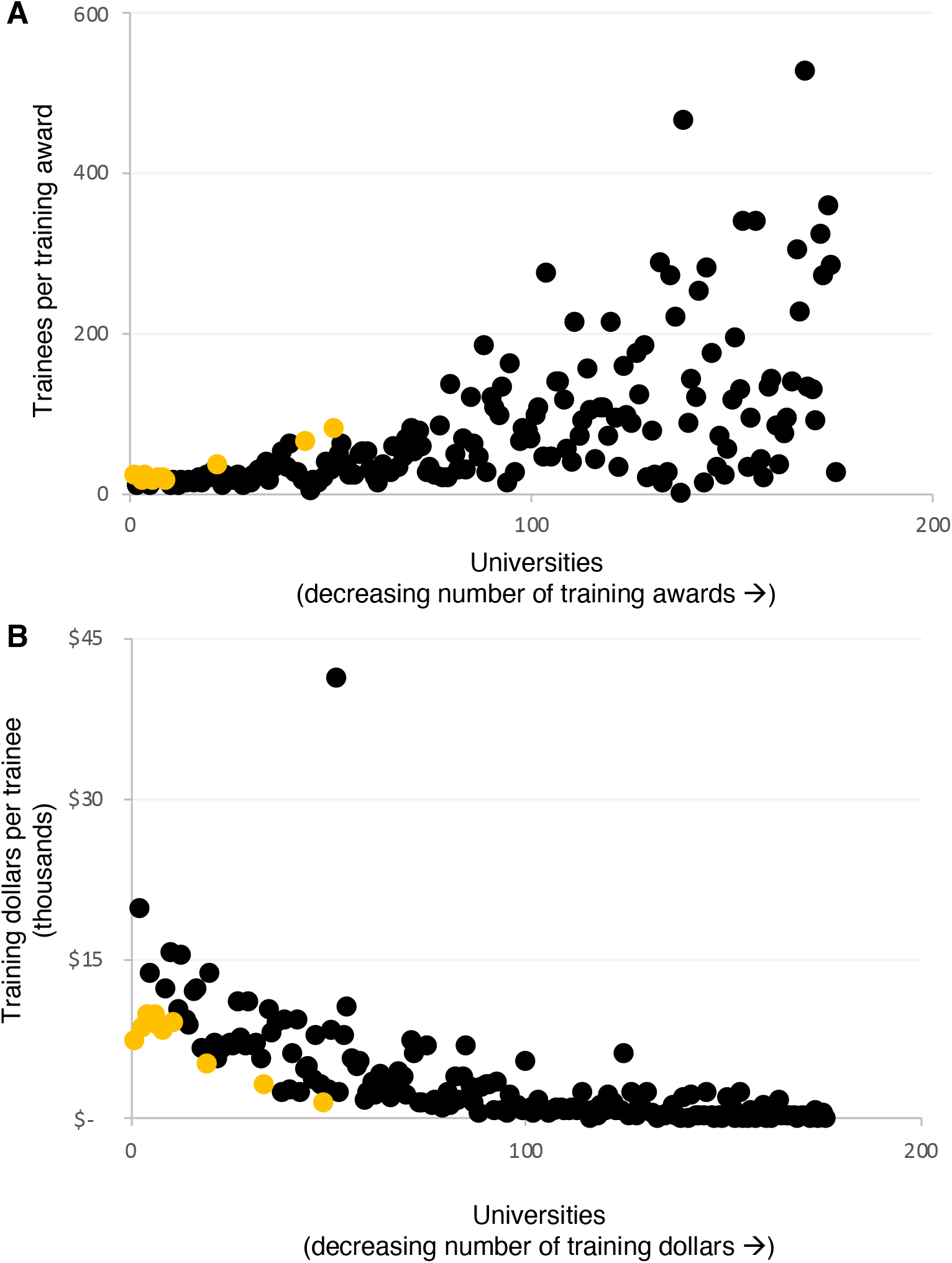
The 2016 training capacity of the biomedical research enterprise. **(A)** The number of trainees per training award at an institution. Institutions are arranged from left to right based on the number of training awards received in 2016. **(B)** The training dollars received by an institution per trainee. Institutions are arranged from left to right based on the number of training dollars received in 2016. Black symbols indicate one institution. Yellow dots indicate the ten institutions discussed in Table 1. For both graphs, N = 176 institutions.

## Discussion

Here I present three Microsoft Excel macros designed to make the NCSES’s GSS data on trainee populations directly relatable to grant award data published by the NIH. With this crosswalk, researchers can mine these databases to better understand how various policies shaped the biomedical research workforce by relating changes in trainee numbers, funding, and awards made, among others. Because the crosswalk uses universities and institutions to relate the databases, it is also possible to determine how specific institutions responded to new policies.

Reports and analyses of the biomedical research enterprise often call for an expansion of publicly available data on the workforce. Some recent progress has been made in universities publishing the career outcomes of their Ph.D. and postdoc alumni (*6-9*). But much more remains. The federal government collects and publishes an incredible amount of information about the research workforce and the money provided to it. By connecting the GSS and NIH databases, researchers can mine new insights from these data without expanding data collection efforts. Furthermore, beyond connecting the GSS and NIH databases, the work here provides a platform to connect other databases that include university names in their data collection efforts. This includes other NCSES databases, like the Survey of Doctoral Recipients, other federal databases, and possibly those of non-governmental organizations. Relating this wide variety of databases will improve our understanding of the research enterprise.

